# M-TUBE: a fabrication-free microfluidic device for large-volume bacterial electroporation requiring minimal assembly

**DOI:** 10.1101/2022.01.14.476275

**Authors:** Po-Hsun Huang, Sijie Chen, Anthony L. Shiver, Rebecca Neal Culver, Kerwyn Casey Huang, Cullen R. Buie

## Abstract

Conventional cuvette-based and microfluidics-based bacterial electroporation approaches have distinct advantages, but they are typically limited to relatively small sample volumes, reducing their utility for applications requiring high throughput such as the generation of mutant libraries. Here, we present a disposable, user-friendly microfluidic electroporation device capable of processing large volume bacterial samples yet requiring minimal device fabrication and straightforward operation. We demonstrate that the proposed device can outperform conventional cuvettes in a range of situations, including across *Escherichia coli* strains with a range of electroporation efficiencies, and we use its large-volume bacterial electroporation capability to generate a library of transposon mutants in the anaerobic gut commensal *Bifidobacterium longum*.

## Main text

One of the key steps in bacterial genetic engineering is the delivery of DNA into cells, which can be realized by mechanical, chemical, or electrical methods^1–3^. Among these methods, electroporation has been the gold standard because it is not cell-type-specific^2^, can deliver molecules of various sizes^4^, and can exhibit relatively high efficiency under optimized conditions^2, 5^. For optimal electric field conditions, genetic material enters cells through reversible pores formed in the cell membrane^6, 7^. Electroporation is typically performed using cuvettes, in an operator-dependent manner that is limited to small batches of volume 1 mL or less. Even with high efficiency, creation of a comprehensive mutant library with hundreds of thousands of mutants^8–10^ for functional-genomics studies can require electroporation of large volumes (tens of milliliters) of saturated bacterial culture, which corresponds to hundreds of cuvette-based electroporation reactions. Performing serial electroporation with manual pipetting is a labor-intensive, time-consuming, and costly process. Moreover, cuvette-based electroporation suffers from issues such as residual volume and joule heating^11, 12^, which affect electroporation efficiency, cell viability, and overall yield.

Performing electroporation in a microfluidic format^11–14^ can remove the need for manual pipetting and improve heat dissipation^11, 14^, thereby increasing electroporation efficiency and cell viability. However, most microfluidic devices involve complicated fabrication processes using PDMS^15–19^, which is an obstacle to widespread adoption, particularly within the microbiology community that would most benefit. Microfluidics-based electroporation devices are also typically limited by the sample volume they can handle. These devices are commonly used for mammalian cells^18, 20^, with just a few examples of applications to bacteria^19, 21^. Several commercial products^22–26^ have demonstrated the potential for scaling up electroporation to throughput of up to ∼100 mL at 8 mL/min^26^, but most have been applied only to mammalian cells and still rely on batch-wise operation^22–26^. Moreover, existing commercial systems require sophisticated electroporation chambers that limit the volume that they can process. Thus, the capabilities of these systems for large-volume bacterial electroporation are yet unproved.

The ideal genetic transformation system would allow for a wide range of sample volumes to accommodate different applications, especially involving the creation of mutant libraries given the low electroporation efficiency of many understudied yet health-relevant bacterial species^10, 27, 28^. A scalable, high-volume electroporation device should be easily assembled by a microbiologist without sophisticated fabrication, compatible with commercially available and common laboratory equipment, and able to process relevant sample volumes in minutes to minimize biological variability. To this end, here we introduce a simple yet powerful <u>M</u>icrofluidic <u>TU</u>bing-based <u>B</u>acterial <u>E</u>lectroporation device (M-TUBE) that enables flexible electroporation of large-volume bacterial samples. M-TUBE facilitates scalable, continuous flow, large-volume bacterial electroporation without the need for micro/nanofabrication, PDMS casting, or 3D printing of microfluidic channels and electrodes.

The M-TUBE device consists of two syringe needles and one plastic tube of a defined length (Fig. 1a). The plastic tubing serves as the microfluidic channel, and the syringe needles serve as the two electrodes, which, when connected to an external high-voltage power supply, establish an electric field across the tubing microchannel. Upon establishing an electric field in the channel, bacterial cells flowing through the channel can be electrotransformed and uptake surrounding genetic material. The syringe needles and plastic tubing used to assemble M-TUBE are commercially and readily available at low cost (<$0.21 per device), and the overall size of an M-TUBE device is similar to that of a conventional cuvette (Fig. 1b). Because syringe needles of standard common formats can be used, M-TUBE can be attached to any commercially available syringe with complementary connectors and can be conveniently interfaced with any syringe pump for sample delivery (Fig. 1c).

**Figure 1:**
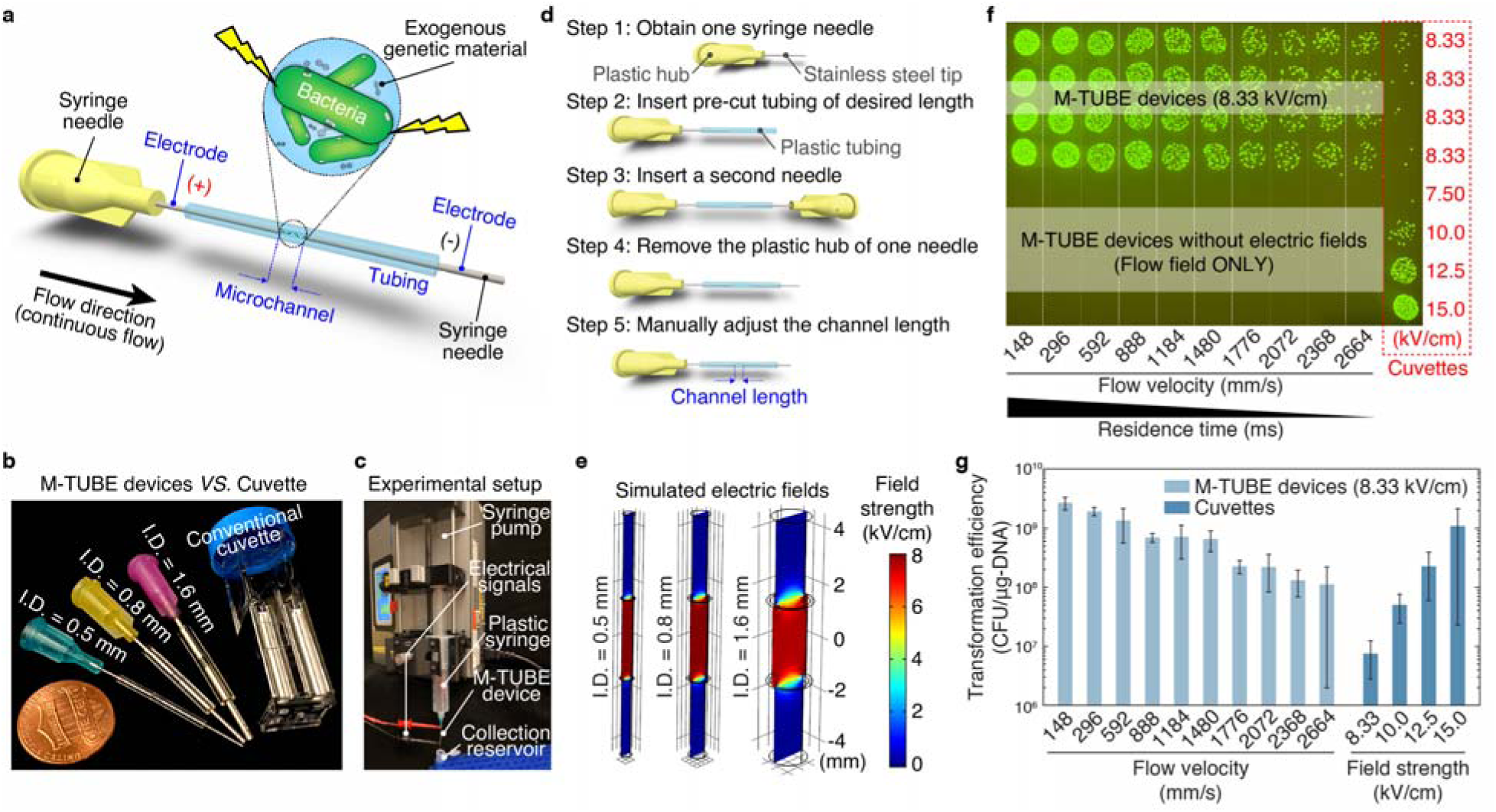
M-TUBE is a fabrication-free, microfluidics tubing-based bacterial electroporation device that is simple to assemble and exhibits higher electroporation efficiency than cuvettes. a) Schematic of the M-TUBE device. The device is composed of two syringe needles and one piece of plastic tubing of pre-defined length. The two syringe needles and plastic tubing serve as the two electrodes and microchannel, respectively. When the two electrodes are connected to an external power supply (or electrical signal generator), an electric field is established within the microchannel, where bacterial electroporation can take place. b) M-TUBE devices with three inner diameters (I.D.) are all similar in size to a conventional cuvette. c) Photograph of the experiment setup when using the M-TUBE device. Since the M-TUBE device is made from standard, commercially available syringe needles and plastic tubing, it can be readily attached to syringe pumps for automated sample delivery, removing the need for manually pipetting samples. d) Detailed breakdown of the protocol for M-TUBE assembly. One device can be completely assembled in 90–120 s. The total cost of parts is currently less than $0.22 and this price could be lowered if parts are bought in bulk. e) Simulations of the electric field established in M-TUBE devices predict similar field strengths irrespective of I.D. f) Spot-dilution assay to quantify viability on selective plates when *E. coli* NEB10β cells were flowed through the device with a plasmid encoding ampicillin resistance and GFP (Table S4) in the presence or absence of an electric field. Transformation was dependent on the electric field. For M-TUBE devices, a voltage of ±2.50 kV (AC field) was applied, which results in an electric field of 8.33 kV/cm. The same batch of cells was used to conduct cuvette-based electroporation as a comparison. g) Comparison of transformation efficiency (colony forming units (CFUs) per μg of DNA) corresponding to the plates in (f). The electroporation efficiency of M-TUBE decreased as the fluid velocity was increased, as expected due to the shorter duration of exposure to the electric field. Regardless of the fluid velocity, the efficiency of M-TUBE was at least one order of magnitude higher than that of cuvettes with the same field strength (8.33 kV/cm). Data represent the average (*n*≥3) and error bars represent 1 standard deviation.

The M-TUBE device can be easily assembled in five steps (Fig. 1d). In brief, device assembly is accomplished by inserting one syringe needle into the plastic tubing cut to a particular length (Methods), and a second syringe needle is inserted into the other end of the tubing. Once both needles are inserted, the length of the channel is manually adjusted to a pre-defined value (Methods) by modifying the gap between the facing ends of the two syringe needles. Assembling a single M-TUBE device requires only 90-120 s (Methods and Video S1), far more convenient than typical fabrication processes for microfluidic devices (usually require several days).

Simulations of the electric field (using COMSOL Multiphysics 5.5) established in the tubing microchannel of M-TUBE (Fig. 1e) indicate that the electric field strength is unaffected by the size of the microchannel (i.e., the tubing inner diameter), assuming that the applied voltage (e.g., 2.50 kV) and distance between the two electrodes (gap, or microchannel length) are held constant. This characteristic enables M-TUBE devices to cover a wider range of sample flow rates without having to adjust the applied voltage to maintain the same field strength. The gap of M-TUBE devices can be easily adjusted without additional assembly, unlike devices that rely on microfabrication, CNC machining, or 3D printing^29^, providing a simple method for adjusting electric field strength of a device. Another beneficial feature is that the residence time within M-TUBEs can be adjusted to control cell exposure to the electric field. Since M-TUBE electroporates bacterial cells in a continuous flow manner, the residence time is dictated by the fluid velocity (or flow rate), such that residence time decreases with an increase in fluid velocity if the gap is fixed (Table S1). These two features, gap length and flow rate, offer users more flexibility in tuning important electroporation parameters such as the electric field strength and the residence time, respectively, which are not always readily tunable in conventional electroporators.

To establish the utility of M-TUBE, optimize its design, and showcase its ability to electrotransform bacterial cells, we used a strain of *E. coli* (NEB10β) with high transformation efficiency. The M-TUBE devices employed for most experiments conducted in this study were comprised of a 500-μm diameter tube and 3-mm gap, and were supplied with a voltage of ±2.50 kV or 5.00 kV_PP_ (peak-to-peak AC signal, square wave), which leads to a field strength of 8.33 kV/cm within the microchannel. Cuvettes with 2-mm gaps were used to perform electroporation at different voltages for as a control. We first confirmed that the flow field (or flow shear stress) along the tube does not by itself lead to genetic transformation. In the absence of an electric field, simply flowing cells through M-TUBE at fluid velocities ranging from 148 mm/s (1.8 mL/min) to 2664 mm/s (32.6 mL/min) did not result in any transformation events (Fig. 1f, bottom). By contrast, once a sufficient electric field was established within M-TUBE, colonies were obtained across the entire range of flow rates tested (Fig. 1f, top), with transformation efficiencies ranging from 10^8^-10^10^ CFUs/μg of DNA (Fig. 1g). A reduction in electroporation efficiency was observed as the fluid velocity was increased.

This trend was expected because the residence time decreases as the flow rate increases, hence cells are exposed to the electric field for a shorter duration at higher flow rates. Despite the lower efficiency at higher flow rates, the overall efficiency obtained using the M-TUBE device was at least one order of magnitude higher than that obtained using cuvettes with the same field strength (8.33 kV/cm). We also note that, compared to cuvettes (typically used at 10-15 kV/cm), M-TUBE was able to produce a comparable efficiency using a lower electric field. The finding that M-TUBE outperforms cuvettes in terms of transformation efficiency may be due to a synergistic effect of the flow field and the electric field^30^.

Given the strong dependence of transformation efficiency on field strength in cuvette-based electroporation, we next evaluated how M-TUBE performs across field strengths. Compared to cuvette-based electroporation at 8.33 kV/cm, regardless of the supplied field strength, M-TUBE exhibited higher transformation efficiencies across the range of flow rates tested (Fig. 2a, left). This finding indicates that M-TUBE can either achieve the same efficiency with lower field strengths or higher efficiency with the same field strength. Moreover, electroporation efficiencies with M-TUBE had a smaller standard deviation than those obtained with cuvette-based electroporation.

**Figure 2:**
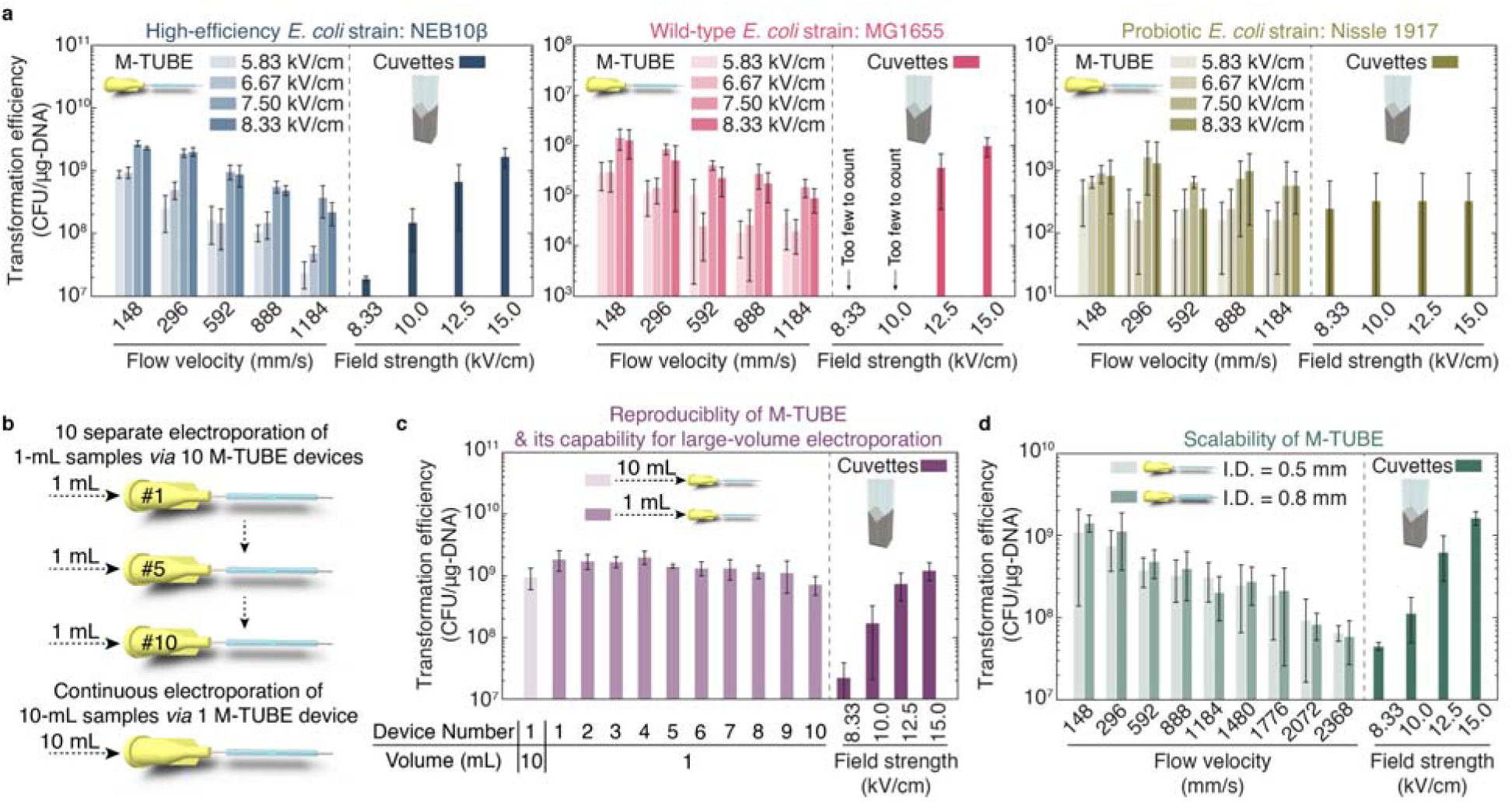
The M-TUBE device exhibits higher efficiency than cuvettes across E. coli strains, is reproducible, and maintains high efficiency across tubing sizes. a) Comparison of M-TUBE device performance when transforming the high-efficiency strain NEB10β, the wild-type strain MG1655, and the probiotic strain Nissle 1917 across voltages and fluid velocities. M-TUBE outperformed cuvettes at an equivalent electric field strength for all strains. Data represent the average (*n*≥3) and error bars represent 1 standard deviation. b) Schematic of the experiment comparing 10 separate 1 mL electroporations and 1 continuous electroporation of a 10 mL sample. c) Transformation efficiency for the experiments in (b) demonstrates that sample volume can be increased without compromising efficiency. Data represent the average (*n*≥3) and error bars represent 1 standard deviation. The same batch of cells was used to conduct cuvette-based electroporation as a comparison. d) Transformation efficiency was similar across 0.5-mm and 0.8-mm diameter M-TUBE devices. For M-TUBE devices, a voltage of ±2.50 kV (AC field) was applied, which results in an electric field of 8.33 kV/cm. Data represent the average (*n*≥3) and error bars represent 1 standard deviation.

Motivated by the successful transformation of *E. coli* NEB10β, M-TUBE was then tested on the wild-type strain *E. coli* MG1655, which typically has lower transformation efficiency than NEB10β. The results show that M-TUBE maintained higher efficiency than cuvettes for MG1655 (Fig. 2, middle). With a field strength of 8.33 kV/cm, M-TUBE yielded efficiencies at least two orders of magnitude higher than cuvettes; even though cuvettes were supplied with a field strength of 10 kV/cm, the number of successfully transformed colonies was too low to reliably enumerate. To further test M-TUBE performance on *E. coli* strains, we used M-TUBE to electroporate the probiotic strain Nissle 1917^27, 28^. While both M-TUBE and cuvettes exhibited much lower electroporation efficiencies for Nissle 1917 compared with MG1655, M-TUBE was comparably efficient to cuvettes and showed slightly better reproducibility (Fig. 2a, right). Moreover, the ability of M-TUBE to process arbitrarily large sample volumes in a continuous fashion means that a desired number of transformed cells of a low-efficiency strain such as Nissle can be obtained with M-TUBE simply by processing a sufficiently large volume. Conversely, using cuvettes for the same goal would be expensive and technically challenging. Overall, M-TUBE showed robust performance across *E. coli* strains with a wide range of electroporation efficiencies, with performance and reproducibility higher than or comparable to cuvette-based electroporation.

Since M-TUBE is hand-assembled, small fluctuations in the microchannel length are inevitable across independently assembled M-TUBE devices (even assembled by the same user). Given that the field strength is defined as the ratio of the applied voltage to the microchannel length, we sought to evaluate if the field strength differs significantly across identical but separately assembled M-TUBE devices, thereby causing variation in electroporation performance for NEB10β cells (Fig. 2b, top). We concurrently carried out electroporation of a large-volume sample (10 mL) to demonstrate the capacity of M-TUBE for high-volume electroporation (Fig. 2b, bottom), from which we were able to determine if there is a substantial difference in transformation efficiency between multiple small volume electroporation experiments and continuous flow large volume electroporation. The variation across 10 M-TUBE devices was insignificant and negligible, and each of the tested devices outperformed cuvettes regardless of the field strength (Fig. 2c), confirming that assembly has negligible impact on the reproducibility of the M-TUBE.

Furthermore, M-TUBE was able to electroporate the entire 10-mL sample at a flow rate of 3.6 mL/min with efficiency higher than or comparable to cuvettes (Fig. 2c) and the transformation efficiency for 10 mL of continuous electroporation was not significantly different from that of 10 separate 1-mL experiments. Continuous electroporation of 10 mL is equivalent to 100 individual 0.1-mL cuvette-based electroporations, for which the configuration of M-TUBE that we tested would shorten the entire electroporation time by two to three orders of magnitude (depending on the flow rate). Put in other terms, M-TUBE can process two to three orders of magnitude more volume of sample in a given period of time compared with cuvettes (Table S2). In terms of cost, M-TUBE is at least 10-fold cheaper than cuvettes (Table S3). Moreover, using M-TUBE for large-volume bacterial electroporation can also circumvent the need for manual pipetting by flowing the electroporated sample directly into recovery medium (Movie S2), thereby decreasing total processing time and potentially improving cell viability and transformation efficiency. Taken together, these features make M-TUBE an ideal candidate for large-volume bacterial electroporation.

Our next goal was to evaluate the ability to scale up the M-TUBE to process even larger volume samples. To this end, the performance of the M-TUBE device with three different inner diameters was compared (500, 800, and 1600 μm, with the size of syringe needles altered accordingly) (Fig. 2d, S1). As long as the gap and the fluid velocity were held fixed, M-TUBE devices with different diameters maintained a high electroporation efficiency for NEB10β cells and outperformed cuvettes. With the same fluid velocity, an M-TUBE device with larger diameter would enable processing larger volumes: with a diameter of 1600 μm, an average fluid velocity of 592 mm/s allows for electroporation of ∼70 mL/min, several orders of magnitude more than what is possible with cuvettes.

These results again demonstrate the capabilities of M-TUBE for large-volume bacterial electroporation, and confirm that M-TUBE can be readily scaled up without compromising efficiency simply by changing the tubing and syringe needles sizes while maintaining fluid velocity.

As a demonstration of the utility of M-TUBE in other organisms, we sought to use the system to generate a set of transposon insertion mutants in a human gut commensal. Many of these organisms are obligate anaerobes and hence require more complex handling during growth, washing, and electroporation. We assembled the M-TUBE electroporation platform inside an anerobic chamber and ran an experiment to generate a small-scale transposon insertion pool in *Bifidobacterium longum* subsp. *longum* NCIMB8809. *B. longum* species are used as probiotics and are actively investigated for their health-promoting effects^31^. To identify optimal electroporation conditions for maximizing transposome delivery, we first electroporated *B. longum* NCIMB8809 cells with the pAM5 plasmid (Fig. 3a, Table S4). As with *E. coli*, M-TUBE plasmid transformation efficiency was comparable to or higher than that of cuvettes for *B. longum* (Fig. 3a). With the optimal electroporation conditions, *B. longum* cells were successfully transformed with *in vitro*-assembled EZ-Tn*5* transposomes, demonstrating its utility both in an anaerobic chamber and for high-throughput transposon mutagenesis (Fig. 3b). Like plasmids, M-TUBE transposome electroporation efficiency was comparable to or higher than that of cuvettes. Thus, we expect M-TUBE should have wide applicability for generation of libraries of thousands of transposon mutants, even in bacterial species with complex growth requirements.

**Figure 3:**
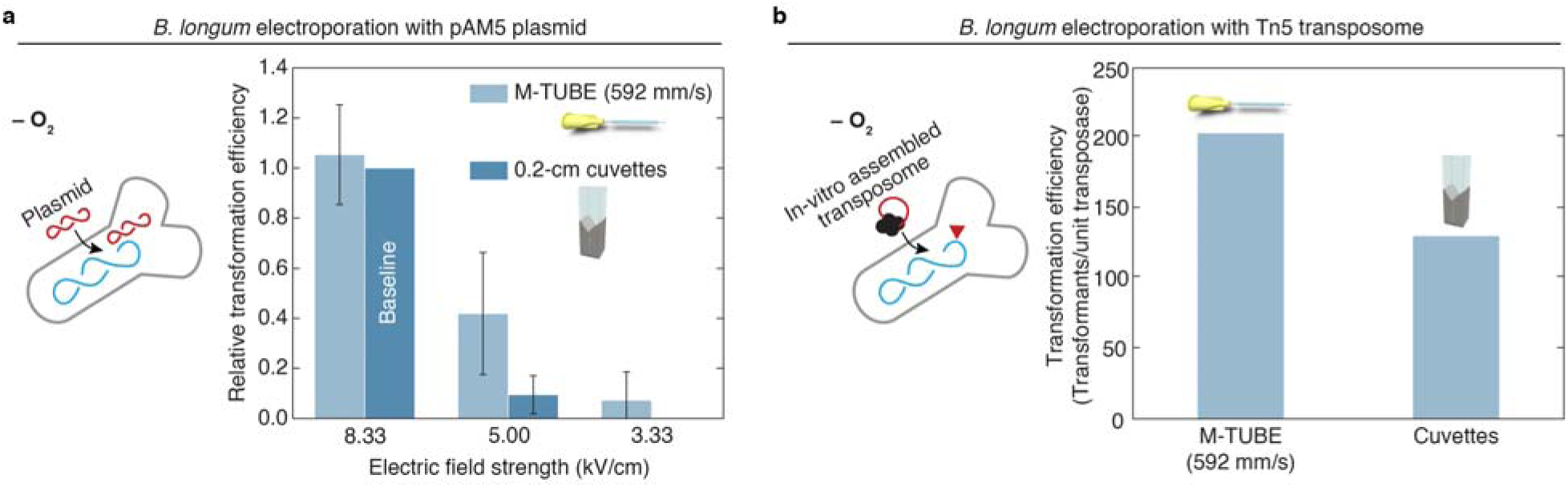
M-TUBE efficiently transforms anaerobic bacteria and enables transposon insertion mutagenesis. a) Comparison of M-TUBE performance during electrotransformation of *B. longum* NCIMB8809 with the plasmid pAM5 at various electric field strengths. For M-TUBE devices, voltages of ±2.50, ±1.50, and ±1.00 kV (AC field) were applied to produce electric fields of 8.33, 5.00, and 3.33 kV/cm, respectively. A fluid velocity of 592 mm/s was used for the M-TUBE device because the ∼5 ms residence time with an M-TUBE inner diameter of 0.5 mm is similar to the time constant observed in cuvette electroporation (5.2-5.6 ms). Data represent the average (*n*≥3) and error bars represent 1 standard deviation. b) Comparison of M-TUBE performance during electrotransformation of *B. longum* NCIMB8809 with Tn5 transposome. For the M-TUBE device, a field strength of 8.33 kV/cm and fluid velocity of 592 mm/s were used, motivated by the results in (a).

Taken together, the disposable, fabrication-free M-TUBE device can process large volumes of bacterial cells with dramatically reduced processing time and effort, without compromising transformation efficiency and cell viability. Due to the simplicity of its fabrication and the wide availability of its components, M-TUBE presents an electroporation strategy that can be immediately implemented in the microbiology community. The flexibility that M-TUBE offers in tuning electroporation conditions such as field strength and residence time make the device a powerful tool for working with hard-to-transform strains. Given the relatively high transformation efficiency compared with cuvettes and its ability to deal with both small and large volumes, M-TUBE has the potential to be a viable alternative to cuvettes and an indispensable tool for applications requiring large volumes such as the creation of mutant libraries.

## Author Contributions

P.-H.H., S.C., A.L.S., R.N.C., K.C.H., and C.R.B. designed the research; P.-H.H., S.C., A.L.S., and R.N.C. performed the research; P.-H.H., A.L.S., and R.N.C. analyzed the data; and all authors wrote and reviewed the manuscript.

## Acknowledgements

We would like to thank members of the Buie and Huang lab for helpful discussions. We thank Douwe van Sinderen for the gift of Bifidobacteria strains and plasmids. We thank Brandon David Fields from the Department of Biological Engineering at MIT for discussion of *E. coli* Nissle 1917 electroporation and for verification of plasmid pAM5 with PCR. The authors acknowledge funding from NIH grant RM1 GM135102 (to K.C.H. and C.R.B.). K.C.H. is a Chan Zuckerberg Biohub Investigator.

## Methods

### Materials

Syringe needles of various gauges (16, 20, or 23) with blunt tips were purchased from CML Supply LLC. Plastic tubing of various diameters were purchased from Cole-Parmer: 0.5-mm inner diameter (ID) (PB-0641901), 0.8-mm ID (EW-07407-70), and 1.6- mm ID (EW-07407-71). Plastic syringes of various volumes with Luer-Lok tips were purchased from Thomas Scientific: 30 mL (BD302832), 20 mL (BD302830), and 10 mL (BD302995). Luria broth (LB) (BD244620) and dehydrated agar (BD214010) were purchased from VWR. MRS broth (BD288130) and Reinforced Clostridial Medium (RCM) (CM0149B) were purchased from Fisher Scientific. Carbenicillin disodium salt (C3416), tetracycline (T7660), L-Cysteine (C7352), α-Lactose monohydrate (L2643), and sucrose (S7903) were purchased from Millipore Sigma.

### Protocol for preparation of an M-TUBE device

An M-TUBE device is assembled from two syringe needles and one piece of plastic tubing with a pre-defined length (Fig. 1d, Video S1). Here, we describe the details of assembly of an M-TUBE device with a microchannel length of 3 mm and a tubing ID of 0.5 mm. First, we cut plastic tubing (50 feet per roll) into 20-mm segments on a cutting mat with metric dimensions. Second, we take two syringe needles of 23 gauge with a tip length of 0.5 in, which has an outer diameter of 0.63 mm that ensures tight fitting between the tubing inner surface and the outer surface of the syringe needle. Next, we insert one of the syringe needles into the tubing and repeatedly rotate back and forth the tubing and/or syringe needle, until the tip of the syringe needle is close to the middle of the tubing and there is also a small portion of the needle for electrical connection that is not inserted into the tubing. We then insert the other syringe needle and rotate back and forth the tubing/syringe needle or the 2^nd^ syringe needle until a gap (*i.e.*, the microchannel length) of a 2-4 mm between the tips of the two syringe needles is established. The gap size can be checked by placing the entire assembly close to a tape measure. After assembling the three components, we remove the plastic hub from either of the syringe needles. Upon removal of the plastic hub, the gap size should then be carefully re-checked with a tape measure, and slight adjustments can be made to establish a gap of 3 mm by gently twisting either needle inward or outward. After this final adjustment, the M-TUBE device is completely assembled.

As discussed above, assembly of one M-TUBE device requires only 60-90 s, hence we typically prepare 50 M-TUBE devices at a time, in ∼1 h. The M-TUBE devices are placed in a Petri dish, which is sterilized in a biosafety cabinet with UV irradiation overnight. After UV sterilization, M-TUBE devices are stored in a -20 °C freezer or refrigerator until just before conducting electroporation experiments, a step similar to the pre-chilling of electroporation cuvettes.

To prepare M-TUBE devices with other tubing sizes, all steps remain unchanged, but it is necessary to ensure that the plastic tubing is assembled with syringe needles that have complementary outer diameters in their tips.

### Culturing and preparation of E. coli strains

Three *E. coli* strains, including NEB10β (New England Biolabs), K-12 MG1655 (Coli Genetic Stock Center, Yale University) and Nissle 1917 (Mutaflor→, Canada), were employed in this study to test the M-TUBE device. The strains, unless otherwise specified, were cultured, harvested, and made electrocompetent using the same conditions. In brief, glycerol stocks were inoculated into two 14-mL cultures tubes containing 6 mL of LB medium and incubated at 37 °C and 250 rpm. The next morning, 5 mL from each overnight culture was inoculated into 245 mL of LB and grown at 37 °C and 200 rpm to an OD_600_ of 0.5-0.7. Note that each set of *E. coli* experiments involved 15-20 mL of electrocompetent cells at OD_600_=10, which required two 250-mL cultures. Each 250 mL culture was divided equally into six 50-mL centrifuge tubes and spun down at 4 °C and 3500 rpm for 10 min using an Allegra 64R centrifuge (Beckman Coulter). The supernatant was discarded and 6 mL of ice-cold 10% glycerol was used to wash and combine the six cell pellets into one suspension. Each 6-mL cell suspension was equally divided into four 2.0-mL microcentrifuge tubes. The eight microcentrifuge tubes generated from the two 250-mL cultures were centrifuged at 4 °C and 8000 rpm for 5 min, the supernatants were discarded, and 1 mL of ice-cold 10% glycerol was used to wash and resuspend the pellet in each of the 8 tube. These washing steps were repeated twice more. Next, all cell pellets were combined into a concentrated suspension using 8 mL of ice-cold 10% glycerol and the cell concentration (typically OD_600_=20-30) was measured using a UV spectrophotometer (UV-1800, Shimadzu). Depending on the measured concentration, a final sample with OD_600_=10 was prepared by adding an appropriate volume of ice-cold 10% glycerol. This sample was placed on ice prior to electroporation. DNA plasmids (Parts Registry K176011)^19^ encoding ampicillin resistance and green fluorescent protein (GFP) were added to this sample at a final concentration of 0.1 ng/μL for NEB10β and MG1655 cultures; for Nissle 1917, a final concentration of 1 ng/ μL was employed so that the number of colony forming units (CFUs) was above the limit of detection. For electroporation, the sample was loaded into a 30-mL plastic syringe (see section on M-TUBE operation).

### B. longum culturing and preparation for M-TUBE electroporation with plasmid DNA

A 5-mL *B. longum* culture was maintained in an anaerobic chamber (Coy) via daily dilution into fresh medium to prepare for electroporation. Briefly, 1 mL of a *B. longum* culture was inoculated into 9 mL of MRS medium in a culture tube, and five additional serially diluted (at 1:10 ratio) cultures were prepared; these six cultures were incubated at 37 °C overnight. The next morning, the optical density of each culture was measured using a spectrometer, and the culture with OD_600_=3-4 was used for subsequent outgrowth. The selected culture was diluted to OD_600_=0.54 in 60-70 mL and grown to OD_600_=1.5-2, after which cells were harvested and made electrocompetent following the same steps described above for *E. coli*. The 60-70 mL were then divided equally into two 50-mL centrifuge tubes and spun down outside the anaerobic chamber at 4 °C and 3500 rpm for 10 min using an Allegra 64R ultracentrifuge (Beckman Coulter). Next, the two 50-mL centrifuge tubes were returned to the anaerobic chamber, the supernatant was discarded, and 5 mL of ice-cold 10% glycerol were used to wash and combine the two cell pellets into one suspension. The 5-mL cell suspension was divided equally into four 2-mL microcentrifuge tubes. The four tubes were centrifuged inside the chamber at room temperature and 10,000 rpm for 2 min using an Eppendorf 5418 microcentrifuge, the supernatants were discarded, and 1 mL of ice-cold 10% glycerol was used to wash and resuspend the pellet in each of the 4 tubes. These washing steps were repeated two more times. Next, all pellets were combined into a concentrated suspension using 5 mL of ice-cold 10% glycerol. Depending on the concentration, the final sample at OD_600_=10 was prepared by adding the appropriate volume of ice-cold 10% glycerol and then placed on ice prior to electroporation. The pAM5 plasmid encoding tetracycline resistance was added to the sample at a final concentration of 2 ng/μL. The mixture of the plasmid DNA with the cells was loaded into a 10-mL plastic syringe for electroporation.

### Transposon mutagenesis of B. longum NCIMB8809

Previous transformation protocols ^32–34^ were combined with minor modifications to prepare electrocompetent cells of *B. longum* NCIMB8809. Briefly, a glycerol stock of *B. longum* NCIMB8809 was recovered for 24 h in 5 mL of MRS broth (MRS media, Difco) at 37 _°_C and passaged overnight (16 h) in 10 mL of MRS in a 10-fold dilution series. The next morning, the incubator temperature was raised to 40 _°_C and one of the overnight cultures in the dilution series was used inoculate 50 mL of MRS (MRS media, HIMEDIA) supplemented with 0.2 M sucrose in a 250-mL Erlenmeyer flask at an initial OD*_600_* (optical density at λ=600 nm) of 0.18, as measured by a 96-well plate reader (Epoch2, BioTek) in a 96-well flat bottom microplate (Grenier Bio-One, Cat. No. 655161) with 200 μL of culture per well. In the dilution series, the overnight culture with the lowest optical density that still provided enough cells to proceed was used to inoculate the next culture. The 50 mL of culture in HIMEDIA-brand MRS was grown to an OD*_600_* of 1.0 and used to inoculate MRS broth reconstituted from individual components— modified with 1% lactose as the sole carbon source, 0.2 M sucrose, and an additional 133 mM NaCl—at an initial OD_600_ of 0.18. This culture was harvested at an OD_600_ of 0.5, pelleted, washed three times with 15% glycerol, and resuspended at an OD_600_ of 6.7 in 15% (v/v) glycerol. To harvest the cells, the culture was moved to a pre-reduced 50 mL conical tube (Fisher Scientific, Cat. No. 06-443-19) on ice, brought out of the anaerobic chamber, centrifuged for 10 min at 3,428*g* (Centrifuge 5920R, Eppendorf), and transferred back into the anaerobic chamber. After cells were harvested, the incubator temperature was lowered back down to 37 °C. Subsequent washes were performed at a volume of 5 mL in 5 mL Eppendorf tubes (Cat. No. 0030122321, Eppendorf) and pelleted with a compatible microcentrifuge (MC-24™ *Touch*, Benchmark Scientific) that had been brought into the chamber, using 2 min 10,000*g* centrifugation steps. Transposomes were assembled *in vitro* by mixing an erythromycin resistance cassette with commercially available EZ-Tn*5* transposase according to manufacturer’s instructions. Transposomes were mixed with competent cells at a concentration of 2U transposase/mL competent cells and electroporated using the M-TUBE device (see below). Electroporated cells were recovered for 2 h at 37 °C, concentrated 10-fold through centrifugation and resuspension in MRS, and plated on RCM-agar plates with 5 μg/mL erythromycin. Colonies were harvested for sequencing after ∼36 h of growth at 37 °C.

### Electroporation of E. coli strains using M-TUBE

The final sample of cells mixed with plasmid DNA was loaded into a plastic syringe, which was mounted on a syringe pump (Legato 210P, KD Scientific) that could be operated horizontally or vertically. To prevent bending of the plastic tubing of the M-TUBE device and to enable convenient collection of the electroporated sample directly into tubes, we typically operate the syringe pump as shown in Fig. 1c. After arranging the pump to operate vertically, an M-TUBE device was attached to the sample-loaded syringe via Luer-Lok connection, and the two syringe-needle electrodes were connected to an external high-voltage power supply system (Fig. S2), which consists of a function generator (3320A, Agilent Technologies), a high-voltage amplifier (623B, Trek Inc.), and an oscilloscope (DSO-X 2022A, Agilent Technologies). Upon confirming a tight connection between the M-TUBE device and the power supply, we pre-filled the M-TUBE microchannel by infusing the cell sample at a relatively low flow rate (typically 250-500 μL/min), to prevent air bubbles and thereby arcing/sparking in M-TUBE, until we visually confirmed that the microchannel was filled with the liquid cell sample. Next, a collection tube (reservoir) was placed underneath the M-TUBE device (Fig. 1c) so that the electroporated sample could be directly and automatically collected. We programmed the pumping parameters including target pumping volume and pumping flow rate, and started flow using the syringe pump at the pre-set flow rate; immediately after starting flow, we started the application of electric signals to the M-TUBE device to initiate electroporation.

As a positive control, the same batch of electrocompetent cells was also electroporated at various field strengths using 0.2-cm electroporation cuvettes (89047-208, VWR). One hundred microliters were pipetted into a pre-chilled electroporation cuvette. Each cuvette was pulsed with an electroporator (MicroPulser™, Bio-Rad) at field strengths including 8.33 kV/cm, 10.0 kV/cm, 12.5 kV/cm, and 15 kV/cm with time constants between 5.0-5.5 ms. Immediately after the application of electric pulses to each cuvette, 900 μL of pre-warmed (∼37 °C) LB recovery medium were added to each cuvette, and the 100-μL electroporated cells was mixed with the 900-μL recovery medium *via* pipetting. We then aspirated as much electroporate sample volume as possible from the cuvette and dispensed it into designated wells on a 96-well deep plate (Fig. S3), along with the electroporated samples from M-TUBE for subsequent recovery at 37 °C for 1 h.

### Electroporation of B. longum via M-TUBE

Most steps for *B. longum* were the same as for *E. coli* described above; the differences are described here. After pre-filling an M-TUBE device with the *B. longum* sample, a 50-mL conical tube (reservoir) containing MRS recovery medium was placed underneath the M-TUBE device (Fig. S4), so that electroporated *B. longum* cells could be directly and automatically flowed into the recovery medium. For *B. longum* electroporation with M-TUBE, one flow rate (7.2 mL/min, or 592 mm/s for the 0.5-mm M-TUBE device) was tested at three field strengths (3.33, 5.00, and 8.33 kV/cm).

As a positive control, the same batch of electrocompetent cells was electroporated at the same three field strengths using 0.2-cm electroporation cuvettes. One hundred microliters of the final cell sample were pipetted into a pre-chilled electroporation cuvette. Each cuvette was pulsed by the electroporator with time constants ranging between 5.4-5.8 ms. Immediately after the application of an electric pulse, 1000 μL of pre-warmed (∼37 °C) LB recovery medium were added to each cuvette and mixed with the cells via pipetting. We then aspirated as much electroporated sample volume as possible from the cuvette and dispensed it into a 1.5-mL microcentrifuge tube.

### Collection, recovery, and evaluation of electroporated E. coli samples

In each set of *<u>E. coli</u>* experiments, a range of flow rates and electric field strengths were tested; for each combination of testing conditions, 1 mL of electroporated sample was collected in a microcentrifuge tube. One hundred microliters of the electroporated sample was aspirated and dispensed into each of four wells of a 96 deep-well plate containing LB recovery medium (Fig. S3). In each 96-well plate, we were able to test 20 combinations of electroporation conditions. After filling all designated wells of the 96- well plate, the plate was incubated in a shaking incubator at 37 °C and 250 rpm for 1 h.

After 1 h of recovery, the 96-well sample plate was placed in a designated position on a liquid handling robot (Janus BioTx Pro Plus, PerkinElmer) for automated serial dilution (Fig. S5): 10X, 100X and 1000X dilution for *E. coli* NEB10β; 10X and 100X dilution for *E. coli* K12 MG1655 or Nissle 1917. Following serial dilution, 5 μL from each well were dispensed onto LB-agar plates containing 50 μg/mL carbenicillin, and the selective plates were incubated overnight at 37 °C. The next morning, each plate was photographed for CFU counting.

### Collection, recovery, and evaluation of electroporated B. longum samples

After electroporating *B. longum* using M-TUBE, 1 mL of cells were flowed directly into 10 mL of MRS recovery medium. *B. longum* samples electroporated by M-TUBE or in cuvettes were incubated at 37 °C for 3 h. Following recovery, 1.1 mL from each M-TUBE or cuvette sample were aspirated and pipetted into separate 1.5-mL microcentrifuge tubes and spun down at 10,000 rpm for 2 min. The supernatants were discarded and 200 μL of MRS medium were added into each 1.5-mL tubes to resuspend the cell pellets. Next, the 200-μL suspension was plated onto RCM-agar plates with 10 μg/mL tetracycline, and the selective plates were incubated at 37 °C for at least 48 h. Following the 48-h incubation, each plate was photographed for CFU counting.

### CFU quantification

Photos of selective plates for electroporation with plasmids were captured using an iPhone 11 (Apple) on a tripod with a remote shutter. The photos were imported to ImageJ (NIH) and CFU.Ai v. 1.1 for enumerating the CFUs. The transformation efficiency was defined as the number of CFUs on selective plates per μg of DNA.

### Data availability

All data used in this manuscript are available upon request from the corresponding author.

