## Supplementary Figures for "M-TUBE: a fabrication-free microfluidic device for large-volume bacterial electroporation requiring minimal assembly"

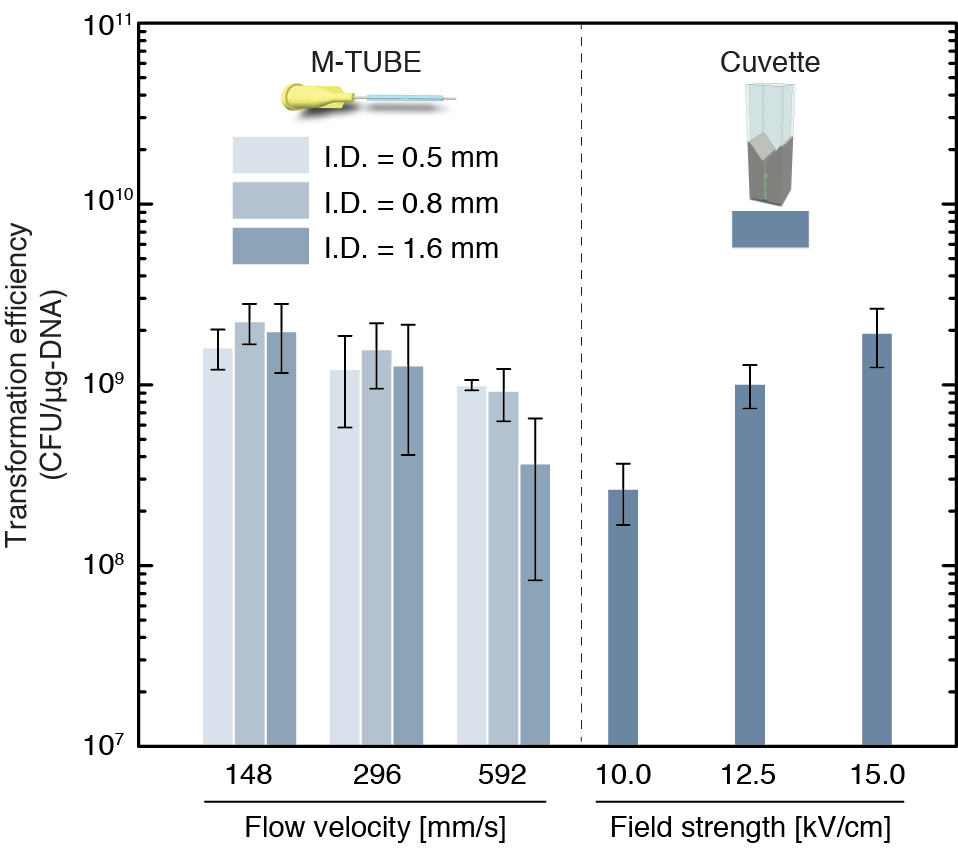

**Figure S1:** **Transformation efficiency is maintained across M-TUBE devices with different diameters.** To further evaluate the scalability of M-TUBE, M-TUBE devices made using plastic tubing with 0.5-mm, 0.8-mm and 1.6-mm inner diameters and compared to conventional cuvettes. A voltage of ±2.50 kV (AC field) was applied to each M-TUBE device, resulting in an electric field of 8.33 kV/cm. The same batch of cells was used to conduct electroporation with 0.2-mm cuvettes and various voltages as a comparison. Data represent the average (*n*≥3) and error bars represent 1 standard deviation.

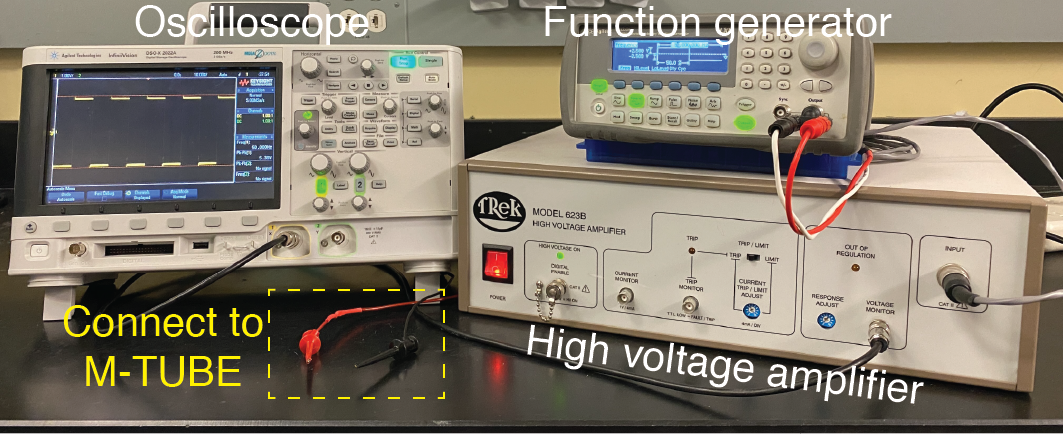

**Figure S2:** **Image of the high-voltage power supply system.** The system is composed of a function generator that allows for waveform programming, a high-voltage amplifier applied to the signal from the function generator, and an oscilloscope that allows for real-time monitoring of the amplified signal.

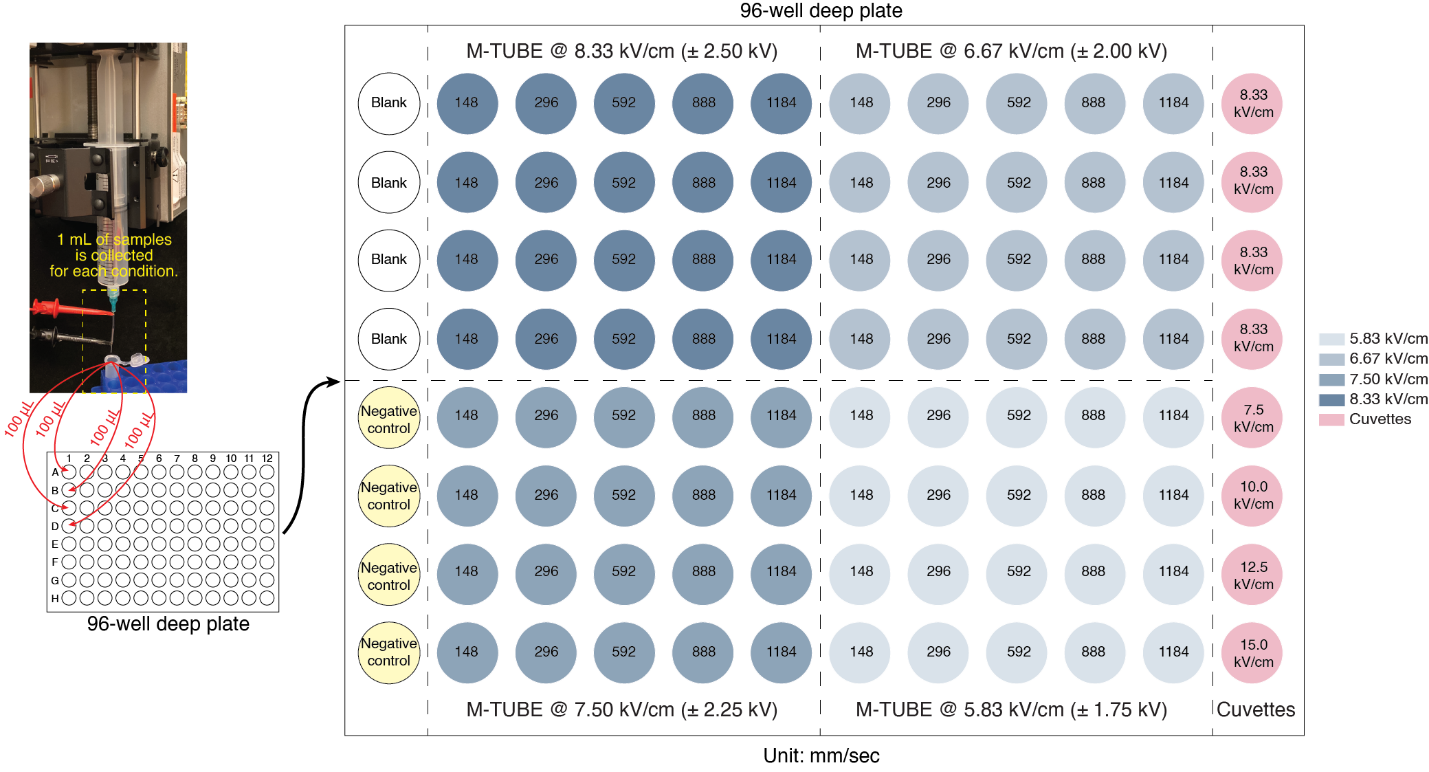

**Figure S3:** **Schematic of the arrangement of electroporation conditions tested in a 96-well deep-well plate.** One milliliter of electroporated cells was collected for each combination of electroporation conditions tested. One hundred microliters were dispensed from each 1-mL sample into each of four designated wells containing 900 µL of LB recovery media. For cuvettes experiments, all of the volume aspirated from each cuvette was dispensed into a well.

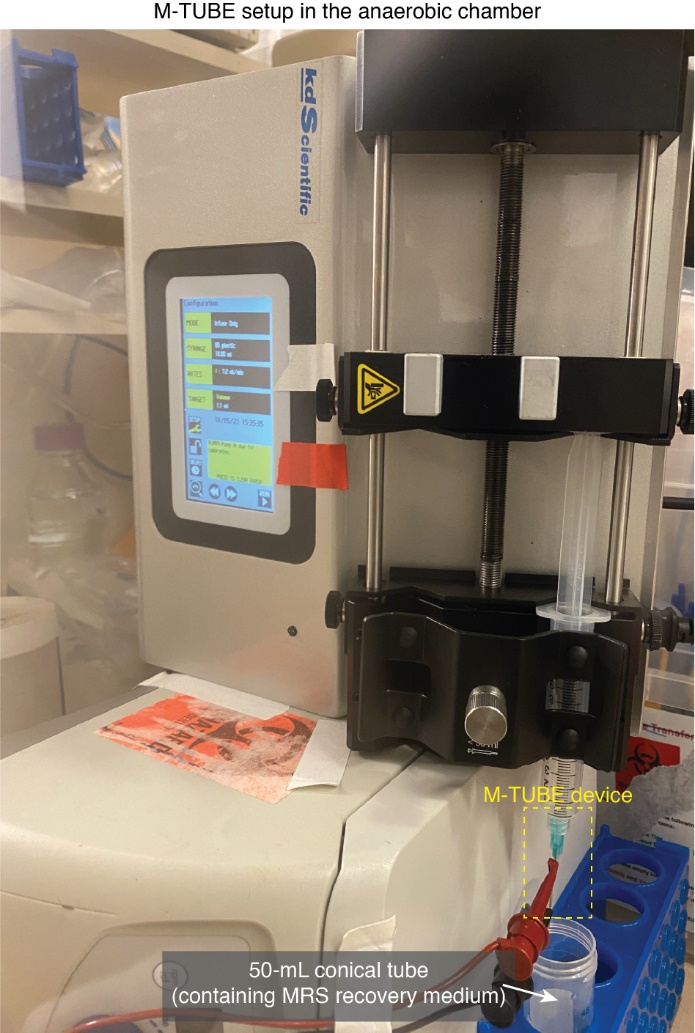

**Figure S4:** **Photograph of M-TUBE set up in an anaerobic chamber.** The M-TUBE device can be easily and conveniently set up in an anaerobic chamber. The photograph also shows that placing a collection tube (reservoir) directly underneath the fluid as it exits the M-TUBE device would enable the direct and automated transfer of electroporated cells into recovery media.

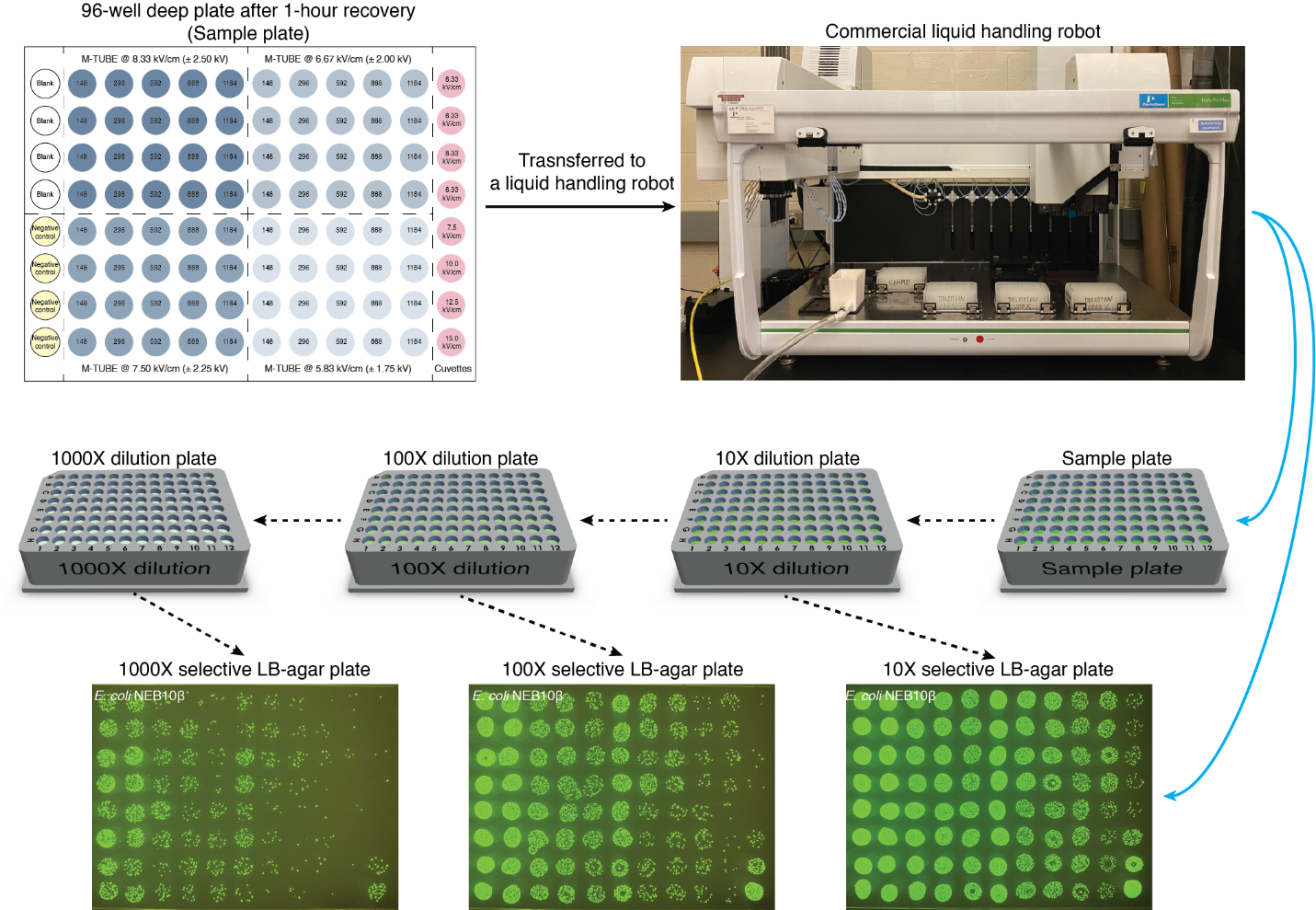

**Figure S5:** **Workflow employing a commercial liquid-handling robot for automated liquid transfer and serial dilution**. After 1 h of recovery, the 96-well deep plate that contains electroporated samples was mounted on a liquid-handling robot. By leveraging the capabilities of the robot, we can use the M-TUBE device to test a wide range of electroporation conditions, each with at least 3-4 technical replicates, while removing the need for extensive manual pipetting for sample transfer, sample dilution, and sample plating. Strain shown is *E. coli* NEB10β.

**Supplementary Tables**

**Table S1:** **Residence time (the duration that cells were exposed to electric fields in M-TUBE devices) as a function of fluid velocities (or flow rates).**

| **M-TUBE device with tubing inner diameter (ID) = 0.5 mm** | | | | | | | | | | |
| --- | --- | --- | --- | --- | --- | --- | --- | --- | --- | --- |
| **Flow velocity**  **(mm/s)** | ~148 | ~296 | ~592 | ~888 | ~1184 | ~1480 | ~1776 | ~2072 | ~2368 | ~2664 |
| **Flow rate**  **(mL/min)** | 1.8 | 3.6 | 7.2 | 10.8 | 14.4 | 18.0 | 21.6 | 25.2 | 28.8 | 32.4 |
| **Residence time**  **(ms)** | **~20.27** | **~10.13** | **~5.07** | **~3.08** | **~2.53** | **~2.03** | **~1.69** | **~1.45** | **~1.27** | **~1.13** |
| **M-TUBE device with tubing ID = 0.8 mm** | | | | | | | | | | |
| **Flow velocity**  **(mm/s)** | ~148 | ~296 | ~592 | ~888 | ~1184 | ~1480 | ~1776 | ~2072 | ~2368 | ~2664 |
| **Flow rate**  **(mL/min)** | 4.4 | 8.8 | 17.6 | 26.4 | 35.2 | 44.0 | 52.8 | 61.6 | 70.4 | 79.2 |
| **Residence time**  **(ms)** | **~20.27** | **~10.13** | **~5.07** | **~3.08** | **~2.53** | **~2.03** | **~1.69** | **~1.45** | **~1.27** | **~1.13** |
| **M-TUBE device with tubing ID = 1.6 mm** | | | | | | | | | | |
| **Flow velocity**  **(mm/s)** | ~148 | ~296 | ~592 | ~888 | ~1184 | ~1480 | ~1776 | ~2072 | ~2368 | ~2664 |
| **Flow rate**  **(mL/min)** | 17.6 | 35.2 | 70.4 | 105.6 | 140.8 | 176.0 | 211.2 | 246.4 | 281.6 | 316.8 |
| **Residence time**  **(ms)** | **~20.27** | **~10.13** | **~5.07** | **~3.08** | **~2.53** | **~2.03** | **~1.69** | **~1.45** | **~1.27** | **~1.13** |

**Table S2:** **Comparison of processing times between conventional cuvettes and M-TUBE devices.** The processing times of cuvettes and M-TUBE devices both scale linearly with sample volume. Across all flow velocities and sample volumes, the M-TUBE device exhibits substantially lower processing time than cuvettes.

| **0.2-cm cuvette (1-1.5 minutes for 100 μL)** | | | | | | | | |
| --- | --- | --- | --- | --- | --- | --- | --- | --- |
| **Processing volume** | | **1 mL** | **5 mL** | **10 mL** | **50 mL** | **100 mL** | **500 mL** | **1,000 mL** |
| Processing time | | 10-15 min | 50-75 min | 1.6-2.5 h | 8-12 h | 16-25 h | 83-125 h | 165-250 h |
| **M-TUBE device (processing time is dependent of flow rates used)** | | | | | | | | |
| **Processing volume** | | **1 mL** | **5 mL** | **10 mL** | **50 mL** | **100 mL** | **500 mL** | **1,000 mL** |
| M-TUBE  (Inner diameter (ID) = **0.5 mm**)  Processing time (min) | 148 mm/s  (1.8 mL/min) | 0.56 | 2.78 | 5.56 | 27.78 | 55.56 | 277.78 | 555.56 |
|  | 296 mm/s  (3.6 mL/min) | 0.28 | 1.39 | 2.78 | 13.89 | 27.78 | 138.89 | 277.78 |
|  | 592 mm/s  (7.2 mL/min) | 0.14 | 0.69 | 1.39 | 6.94 | 13.89 | 69.44 | 138.89 |
|  | 888 mm/s  (10.8 mL/min) | 0.09 | 0.46 | 0.93 | 4.63 | 9.26 | 46.30 | 92.59 |
|  | 1184 mm/s  (14.4 mL/min) | 0.07 | 0.35 | 0.69 | 3.47 | 6.94 | 34.72 | 69.44 |
| M-TUBE  (ID = **0.8 mm**)  Processing time (min) | 148 mm/s  (4.4 mL/min) | 0.23 | 1.14 | 2.27 | 11.36 | 22.73 | 113.64 | 227.27 |
|  | 296 mm/s  (8.8 mL/min) | 0.11 | 0.57 | 1.14 | 5.68 | 11.36 | 56.82 | 113.64 |
|  | 592 mm/s  (17.6 mL/min) | 0.06 | 0.28 | 0.57 | 2.84 | 5.68 | 28.41 | 56.82 |
|  | 888 mm/s  (26.4 mL/min) | 0.04 | 0.19 | 0.38 | 1.89 | 3.79 | 18.94 | 37.88 |
|  | 1184 mm/s  (35.2 mL/min) | 0.03 | 0.14 | 0.28 | 1.42 | 2.84 | 14.20 | 28.41 |
| M-TUBE  (ID = **1.6 mm**)  Processing time (min) | 148 mm/s  (17.6 mL/min) | 0.06 | 0.28 | 0.57 | 2.84 | 5.68 | 28.41 | 56.82 |
|  | 296 mm/s  (35.2 mL/min) | 0.03 | 0.14 | 0.28 | 1.42 | 2.84 | 14.20 | 28.41 |
|  | 592 mm/s  (70.4 mL/min) | 0.01 | 0.07 | 0.14 | 0.71 | 1.42 | 7.10 | 14.20 |
|  | 888 mm/s  (105.6 mL/min) | 0.01 | 0.05 | 0.09 | 0.47 | 0.95 | 4.73 | 9.47 |
|  | 1184 mm/s  (140.8 mL/min) | 0.01 | 0.04 | 0.07 | 0.36 | 0.71 | 3.55 | 7.10 |

**Table S3:** **Comparison of** **costs for assembly of one M-TUBE device versus cuvettes per unit processing volume.**

| **Parts cost for M-TUBE devices and conventional cuvettes** | | | | | |
| --- | --- | --- | --- | --- | --- |
| **Parts for M-TUBE** | | **Quantity** | | **Bulk Price (USD)** | **Note** |
| Syringe needle | | 1000 pieces | | <$101.9 | (<$0.10 per piece) |
| Plastic tubing | | 3048 cm | | <$58.9 | (<$0.02 per cm) |
|  | |  | |  | **(One M-TUBE device costs <$0.22)** |
| **Parts for cuvettes** | | **Quantity** | | **Bulk Price (USD)** | **Note** |
| 0.2-cm cuvette from VWR | | 50 pieces | | $111.38 | ($2.23 per cuvette) |
| 0.2-cm cuvette from BIO-RAD | | 50 pieces | | $117.75 | ($2.36 per cuvette) |
| **Cost to electroporate a unit volume sample (parts only)** | | | | | |
| **Processing volume** | **0.2-cm cuvette from VWR** | | **0.2-cm cuvette from BIO-RAD** | | **M-TUBE device** |
| 1 mL | $22.30 | | $23.60 | | <$0.22 |
| 5 mL | $111.38 | | $117.75 | | <$0.22 |
| 10 mL | $222.76 | | $235.50 | | <$0.22 |
| 50 mL | $1,113.80 | | $1,177.50 | | <$0.22 |
| 100 mL | $2,227.60 | | $2,355.00 | | <$0.22 |
| 500 mL | $11,138.00 | | $11,775.00 | | <$0.22 |
| 1000 mL | $22,276.00 | | $23,550.00 | | <$0.22 |

**Table S4: Strains, plasmids, and oligos used in this study.**

| **Strain** | | **Source** | |
| --- | --- | --- | --- |
| *E. coli* NEB10β | | New England Biolabs | |
| *E. coli* K-12 MG1655 | | Yale Coli Genetics Stock Center | |
| *E. coli* Nissle 1917 | | Mutaflor® | |
| *Bifidobacterium longum* subsp. *longum* NCIMB8809 | | Gift from Douwe van Sinderen, University College Cork, Ireland | |
| **Plasmid** | **Reference** | **Source** | |
| pCon1.00 (J23100)->RBS+GFP+T | <http://parts.igem.org/Part:BBa_K176011> | iGEM | |
| pAM5 | ^35^ | Gift from Douwe van Sinderen, University College Cork, Ireland | |
| **Oligo** | **Sequence** | **Description** | **Source** |
| SHGA080 | /5Phos/CTGTCTCTTATACACATCTATTTATGTTACAGTAATATTGACTTCGACACC | Forward primer for generating randomly barcoded erm^R^ transposon | This study |
| SHGA306 | /5Phos/CTGTCTCTTATACACATCT GTCGACCTGCAGCGTACG NNNNNNNNNNNNNNNNNNNN AGAGACCTCGTGGACATC TTACACATTATTCCGGTGATAGGGC | Reverse primer for generating randomly barcoded erm^R^ transposon | This study |
